# A Novel Simple ImmunoAssay for Quantification of Blood Anti-NMDAR1 Autoantibodies

**DOI:** 10.1101/2024.04.11.589086

**Authors:** Melonie Vaughn, Susan Powell, Victoria Risbrough, Xianjin Zhou

**Author notes:** To whom correspondence should be addressed: Department of Psychiatry, University of California San Diego, La Jolla, USA.

## Abstract

High titers of anti-NMDAR1 autoantibodies in human brain cause anti-NMDAR1 encephalitis, a rare disease that displays a variety of psychiatric symptoms and neurological symptoms. Currently, immunohistochemical staining and cell-based assays are the standard methods for detection and semi-quantification of the anti-NMDAR1 autoantibodies. Low titers of blood circulating anti-NMDAR1 autoantibodies have been reported in a significant subset of the general human population. However, detection and quantification of these low titers of blood circulating anti-NMDAR1 autoantibodies are problematic because of high non-specific background from less diluted serum/plasma. Development of a new method to quantify these low titers of blood anti-NMDAR1 autoantibodies is necessary to understand their potential impacts on psychiatric symptoms and cognition. Based on our previous One-Step assay, we report the development of a novel simple immunoassay to quantify cross-species blood anti-NMDAR1 autoantibodies, and its validation with immunohistochemistry and cell-based assays in both humans and mice.

## Introduction

High titers of anti-NMDAR1 autoantibodies in the brain can cause anti-NMDAR1 encephalitis, a rare disease that displays a variety of psychiatric symptoms and neurological symptoms (Dalmau, 2016). Detection and semi-quantification of anti-NMDAR1 autoantibodies for investigation and diagnosis are most commonly performed via immunohistochemical staining of rodent brains or the cell-based assay (Gresa-Arribas et al., 2014). The potential for low titers of blood anti-NMDAR1 autoantibodies to contribute to the development of psychiatric symptoms has attracted many studies on blood circulating anti-NMDAR1 autoantibodies in the general human population and psychiatric patients (Castillo-Gomez et al., 2017; Hammer et al., 2014; Jezequel et al., 2017; Pan et al., 2019) and in mice (Yue et al., 2021). However, these semi-quantifications of low titers of human blood circulating anti-NMDAR1 autoantibodies were conducted solely using the cell-based assays. We previously found that in human blood, detecting low titers of anti-NMDAR1 autoantibodies using the cell-based assays suffers from high non-specific background (Zhou, 2021a). Additionally, these semi-quantitative methods are subjective in nature. Therefore, developing an objective method to quantify low titers of blood anti-NMDAR1 autoantibodies is important to advance our understanding of their potential impacts on psychiatric symptoms and cognition. We previously reported a GFP-based quick immunoassay to detect antibodies (Yue et al., 2021; Zhou, 2021b). Using the same strategy, we report the development of a *Gaussia* luciferase-based immunoassay to quantify and compare the levels of blood anti-NMDAR1 autoantibodies between mice and humans as well as the validation of the luciferase-based immunoassay with the GFP-based assay, immunohistochemistry, and cell-based assays.

## Materials and methods

### Expression of NMDAR1-luciferase fusion protein

Human NMDAR1 protein sequence was from NCBI reference sequence database (Accession NP_0155661.1). The ligand binding domain (LBD) of NMDAR1 was synthesized with a linker GSGSG according to literature (Furukawa and Gouaux, 2003) and cloned into pET-21d vector. BL21(DE3)pLysS competent *E coli* cells were purchased from EMD (cat. 70236-3) for transformation of the plasmids. The fusion proteins of the NMDAR1 LBD with either GFP or *Gaussia* luciferase were expressed in *E coli* and purified with HisPur Ni-NTA resins (Zhou, 2021b) before solubilized as described (Waldo et al., 1999). Successful folding of the fusion proteins was examined with either GFP fluorescence or activities of *Gaussia* luciferase.

### Human plasma samples

Human plasma samples were leveraged from an existing sample repository with approval from the San Diego Veterans Affairs Healthcare Services (IRB Protocol H180112). Plasma samples were collected with BD Vacutainer™ Plastic Blood Collection Tubes with Lithium Heparin after written consent was obtained. Samples were stored in -80 freezers until use. Samples are from healthy males aged 19 to 23 years old.

### ImmunoAssay

Protein A/G/L was purchased from Novus Biologicals (NBP2-34985) and diluted to 1ug/ul with antibody diluent solution (DAKO, S080983-2). Two microliters of mouse or human serum/plasma was incubated with NMDAR1-GLUC and 2 ul of protein A/G/L in 1X PBS, 0.25M NaCl, 1% Triton X100. The mixture was then incubated for 1.5 hours at room temperature. After adding 200 ul of 1XPBS, 0.1% Tween 20 for washing, the pellet was collected by centrifugation at 3220 rpm for 10 minutes. The pellet was washed again with 200 ul of 1XPBS. After the washing, the pellet was suspended in 10 ul of 1XPBS. 20 ul of *Gaussia* luciferase substrate (ThermoFisher, cat. 16160; Pierce™ Gaussia Luciferase Glow Assay Kit) was then added to the suspended pellet. *Gaussia* luciferase activities were stabilized for 10 minutes at room temperature. Luciferase activities (RLU) were measured with Greiner 96-well Flat Bottom Black Polystyrene plate (Cat. No.: 655097) on Tecan infinite 200Pro.

### Mouse strain

C57BL/6J mice were purchased from Jackson Labs (Bar Harbor, ME) at 8-weeks-old and active immunization was conducted after a week acclimation as described in our previous studies (Yue et al., 2021). All testing procedures were approved by University of California San Diego and San Diego Veterans Affairs Healthcare Services Animal Care and Use Committee prior to the onset of the experiments. Mice were maintained in American Association for Accreditation of Laboratory Animal Care approved animal facilities at the local Veteran’s Administration Hospital or UCSD campus. These facilities meet all Federal and State requirements for animal care.

### Active immunization

P2 peptide antigens of mouse NMDAR1 were synthesized by *Biomatik*. The peptide immunization was conducted as described in our previous studies (Yue et al., 2021). In brief, the P2 peptide was dissolved in PBS at a concentration of 4 mg/ml and mixed with an equal volume of Complete Freund’s Adjuvant to generate a thick emulsion. Mice were injected with 100 ul of emulsion subcutaneously. Control mice were injected with CFA emulsion without the P2 peptide. Two months after immunization, a few microliters of blood were collected from mouse tail vein for analysis of anti-NMDAR1 autoantibodies.

### Immunohistochemistry

Immunohistochemistry (IHC) was conducted as previously described (Ji et al., 2013; Kim et al., 2012) to semi-quantify the levels of blood anti-NMDAR1 autoantibodies in individual mice. Wildtype mouse brain paraffin sections were used for IHC analysis with a dilution of mouse serum at 1:200 with antibody diluent solution (DAKO, S080983-2). Mouse anti-NMDAR1 monoclonal antibody (BD, cat. 556308) was used as the positive control with a dilution at 1:40,000. ImmPRESS peroxidase-micropolymer conjugated horse anti-mouse IgG (H + L) (Vector Labs, MP-7402) was used as the secondary antibody. Biotinylated goat anti-human IgG (H + L) secondary antibody was used for detection of human plasma anti-NMDAR1 autoantibodies. Chromogenic reaction was conducted with ImmPACT NovaRED Peroxidase Substrate (Vector Labs, SK-4805). Slides were mounted with Cytoseal 60 mounting medium (Richard-Allan Scientific, 8310-16). *Image J* was used to measure differential optical intensities between hippocampal CA1 *st oriens* and *corpus callosum* to semi-quantify the levels of anti-NMDAR1 autoantibodies.

### Cell-based assay

Immunofluorescence analysis of mouse and human anti-NMDAR1 autoantibodies were conducted using Euroimmun BIOCHIP with a positive control of human anti-NMDAR1 autoantibody as described in our previous studies (Yue et al., 2021) . Mouse serum and human plasma were diluted at 1:10 and 1:50 respectively with DAKO antibody diluent solution. Goat anti-Human IgG (H+L) Fluorescein (Vector Lab, FI-3000) and goat anti-mouse IgG (H+L) Alexa Fluor 568 (Invitrogen, A11004) were diluted at 1:1,000 as the secondary antibodies for immunofluorescence staining. Flurorescence staining was examined using microscope EVOS FL (ThermoFisher, Scientific).

## Results

We previously developed the One-Step GFP-based assay that enables instant visualization of antibody using protein AGL to aggregate all isotypes of antibodies during antibody binding GFP-labelled antigens (Yue et al., 2021; Zhou, 2021b). To quantify the levels of blood anti-NMDAR1 autoantibodies recognizing NMDAR1 ligand binding domain, we fused the ligand binding domain of human NMDAR1 (Furukawa and Gouaux, 2003) with *Gaussia* luciferase **(Figure 1A)**. Mouse anti-NMDAR1 autoantibodies generated in our previous studies (Yue et al., 2021) were detected with either the NMDAR1-GFP **(Figure 1B)** or quantified with the NMDAR1 fused *Gaussia* luciferase **(Figure 1C)**. We observed negligible non-specific background in either assay, supporting feasibility for developing a luciferase-based assay to quantify the levels of the blood anti-NMDAR1 autoantibodies.

**Figure 1.**
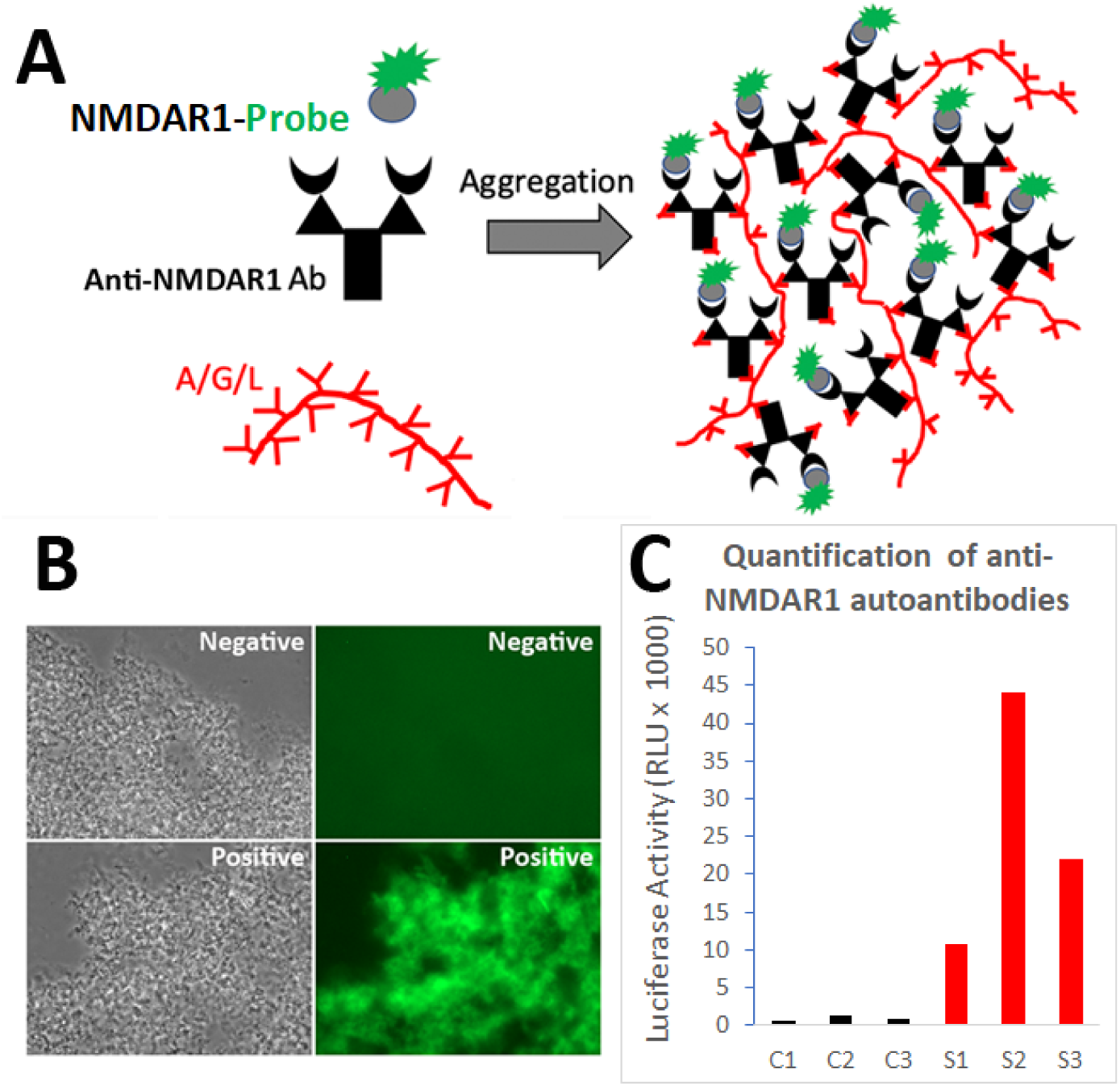
Development of a luciferase-based immunoassay to quantify blood anti-NMDAR1 autoantibodies. **(A)** Strategy for the One-Step Immunoassay (Yue et al., 2021; Zhou, 2021b). **(B)** One-Step immunoassay when the probe is GFP. Blood from negative control mice and mice carrying the anti-NMDAR1 P2 antibodies are shown. All isotypes of blood antibodies are aggregated by protein A/G/L (gray image). Anti-NMDAR1 P2 autoantibodies within the aggregate from a positive mouse bind NMDAR1-GFP and emit strong green fluorescence. Aggregated antibodies from negative control mouse blood showed little background. **(C)** Luciferase-based immunoassay when the GFP probe is replaced by *Gaussia* luciferase. Quantification of the levels of blood anti-NMDAR1 autoantibodies in positive mice (S1-S3) and negative control mice (C1-C3).

To validate the luciferase-based immunoassay, we conducted correlation studies of mouse blood anti-NMDAR1 autoantibodies with immunohistochemistry. A cohort of mice were immunized with the NMDAR1 P2 antigens as shown in our previous studies (Yue et al., 2021). Immunized mice developed anti-NMDAR1 autoantibodies, as confirmed by immunohistochemical staining **(Figure 2A)**. For semi-quantification by immunohistochemistry, optical intensities of CA1 staining (CA1 *st oriens*), after subtracting the background intensity from *corpus callosum*, were used to quantify the relative levels of the blood anti-NMDAR1 autoantibodies for individual mice. The levels of the blood anti-NMDAR1 autoantibodies of individual mice were also quantified using the luciferase-based immunoassay. A high correlation (R^2^=0.97) was observed between the levels of blood anti-NMDAR1 autoantibodies quantified with immunohistochemistry and the luciferase-based assay **(Figure 2B)**. However, ceiling effects seemed to impact the highest levels of anti-NMDAR1 autoantibodies in the immunohistochemistry assay. This finding suggests that more serum dilutions are needed to accurately quantify the high levels of anti-NMDAR1 autoantibodies to avoid color saturation in chromogenic immunohistochemistry. In contrast, activities of *Gaussia* luciferase have a linear range from a thousand to a million relative light units (RLU). Such ceiling effects are not present in the luciferase-based assay; and no dilutions are needed to achieve accurate quantification of high or low levels of the anti-NMDAR1 autoantibodies.

**Figure 2.**
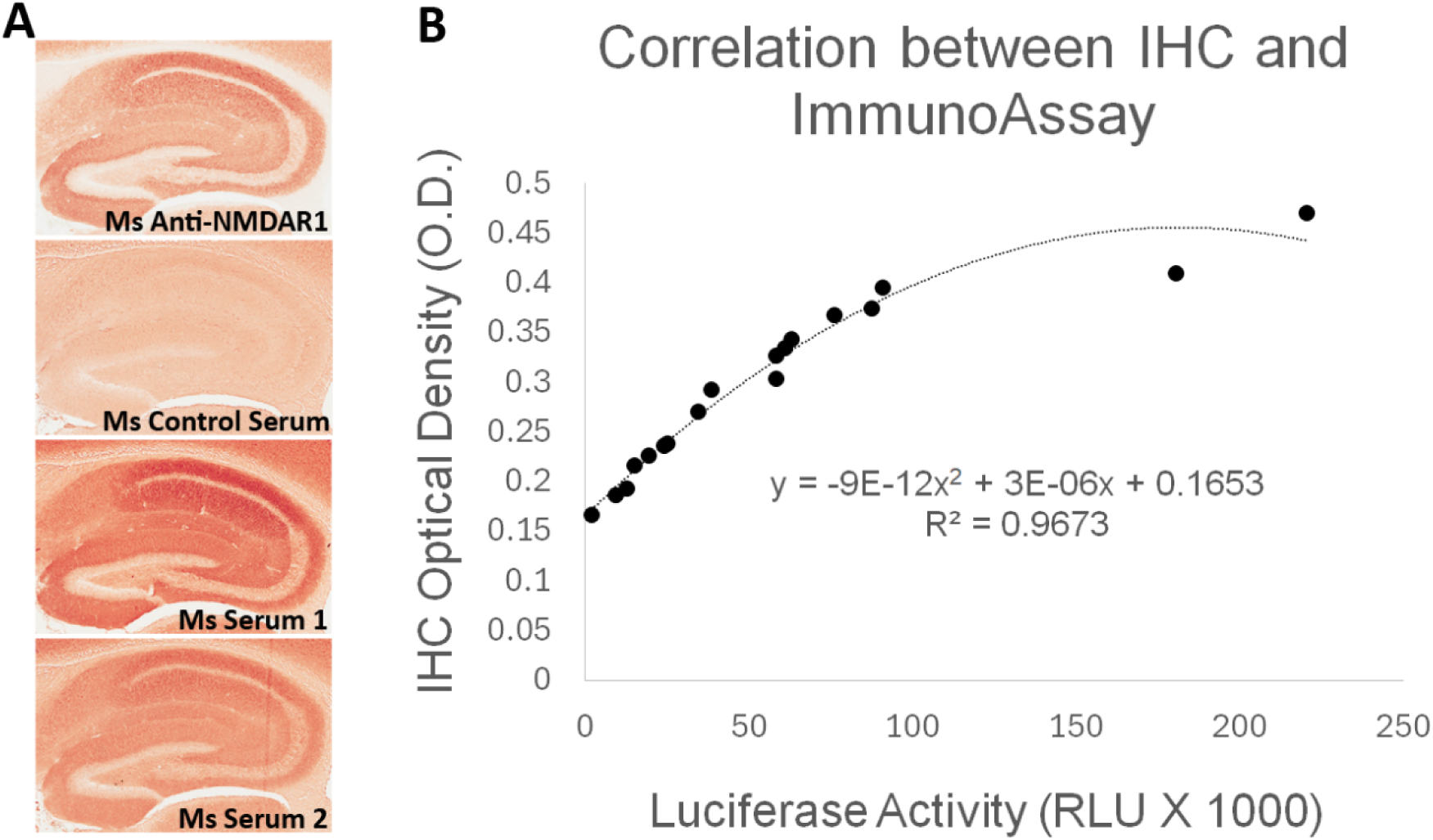
Correlation analysis between quantification by the luciferase-based immunoassay and semi-quantification by immunohistochemistry (IHC). **(A)** Wildtype mouse paraffin hippocampal sections were used for immunohistochemical analysis of anti-NMDAR1 autoantibodies. Sera of immunized mice were diluted at 1:200 for the IHC analysis. Positive control (Ms Anti-NMDAR1): a commercial mouse anti-NMDAR1 monoclonal antibody (1:40,000 dilution). Negative control (Ms Control Serum): mouse immunized with CFA only. Test mice (Ms Serum 1 and 2): representatives from 18 mice immunized with CFA and NMDAR1 P2 antigens. Optical intensities of hippocampal CA1 *st oriens* and *corpus callosum* were quantified with Image J and their differences were used as the relative levels of anti-NMDAR1 autoantibodies in individual mice. **(B)** Anti-NMDAR1 autoantibodies from 18 mouse serum samples were quantified by the luciferase-based immunoassay. A correlation between the IHC semi-quantification and the luciferase quantification of blood anti-NMDAR1 autoantibodies was shown for these 18 serum samples. A strong correlation (R^2^=0.97) was observed between these two methods.

Since the protein A/G/L binds all isotypes of antibodies in mammals, we conducted comparative analyses of the levels of blood circulating anti-NMDAR1 autoantibodies between mice and humans using our luciferase-based immunoassay **(Figure 3)**. All mice immunized with NMDAR1 P2 peptide antigen (red dots) except one have higher levels of anti-NMDAR1 autoantibodies than the background level from the negative control mice (black dots). Human plasma (blue dots) samples were taken from 143 young healthy males with ages ranging from 19 to 23 years old. A higher background level of natural anti-NMDAR1 autoantibodies was observed in human plasma than in mice. The physiological significance of this background difference is unknown, and the difference may be partly attributed to higher concentrations of overall blood antibodies in humans than in mice. Inter Quartile Range (IQR) was used to identify statistical outliers (Q3+1.5*IQR); and more than 10% of the human subjects were the outliers with higher levels of anti-NMDAR1 autoantibodies, supporting previously reported qualitative findings that a significant subset of the general human population carries anti-NMDAR1 autoantibodies in blood.

**Figure 3.**
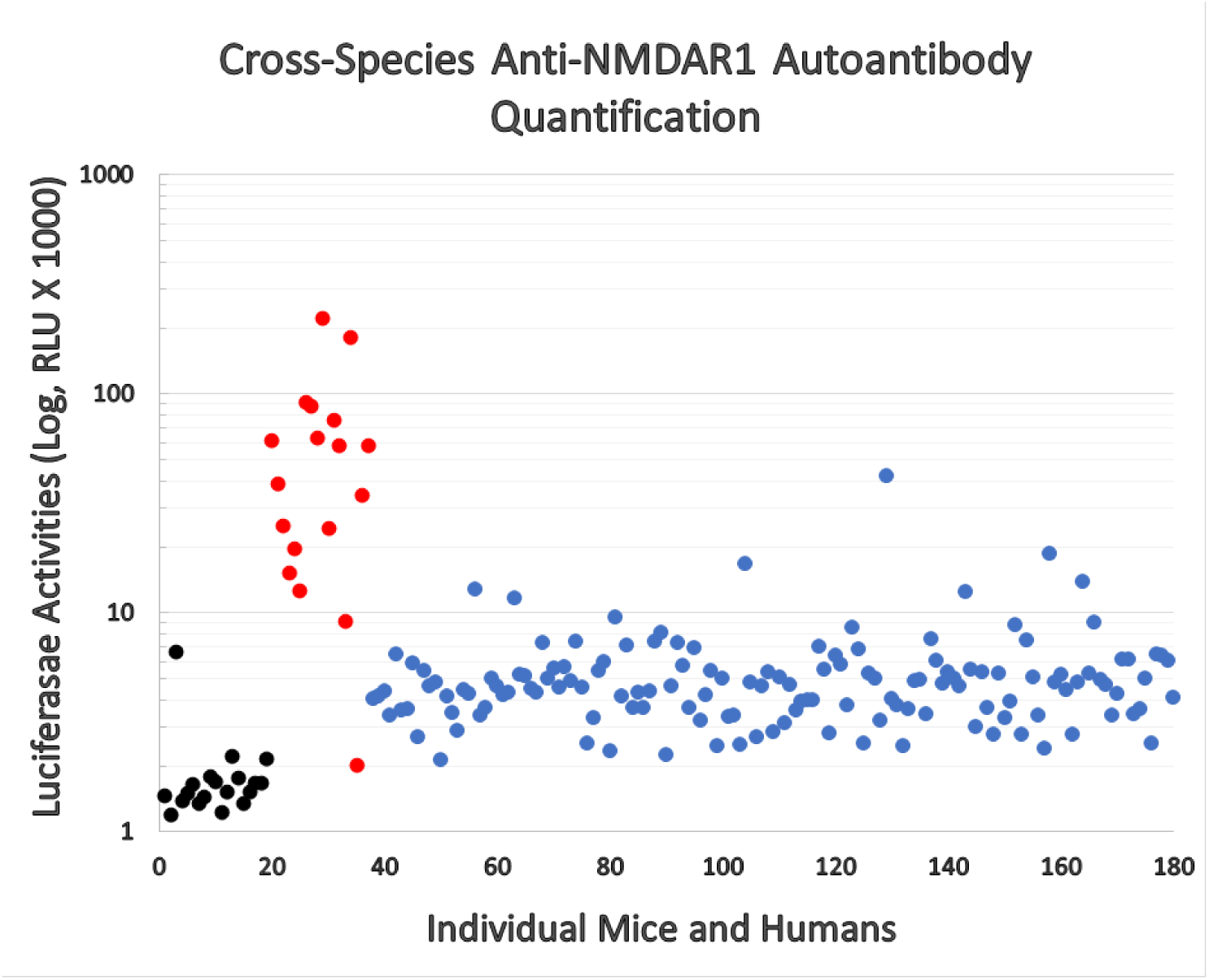
Cross-species quantification and direct comparison of blood anti-NMDAR1 autoantibodies between mice and humans. Anti-NMDAR1 autoantibodies were quantified in the blood of 143 human subjects (blue dots), negative control mice (black dots), and mice immunized with NMDAR1 P2 antigen (red dots). Activities of *Gaussia* luciferase (RLU, X1000) are used as the levels of anti-NMDAR1 autoantibodies (Log10 scale). A significant portion of the general human population carries natural anti-NMDAR1 autoantibodies in blood.

Binding of plasma antibodies to the *Gaussia* luciferase may potentially be detected as false positives of anti-NMDAR1 autoantibodies by the NMDAR1-*Gaussia* luciferase probe. Therefore, we re-examined the quantified samples that carry high levels of anti-NMDAR1 autoantibodies with a different NMDAR1-GFP fusion probe. The top 20 plasma samples carrying the highest levels of anti-NMDAR1 autoantibodies were also positive for the GFP-based assay, ruling out the presence of the antibodies significantly cross-reacting with *Gaussia* luciferase in the 143 human plasma samples.

After quantifying > 300 human plasma samples, we found no significant non-specific binding to *Gaussia* luciferase. Out of these 143 samples, the human plasma carrying the highest levels of anti-NMDAR1 autoantibodies emit strong green fluorescence after protein A/G/L aggregation in contrast to the plasma with basal levels of anti-NMDAR1 autoantibodies **(Figure 4)**. Different NMDAR1 fusion probes with either the GFP or the luciferase provide a cross validation for analysis of anti-NMDAR1 autoantibodies.

**Figure 4.**
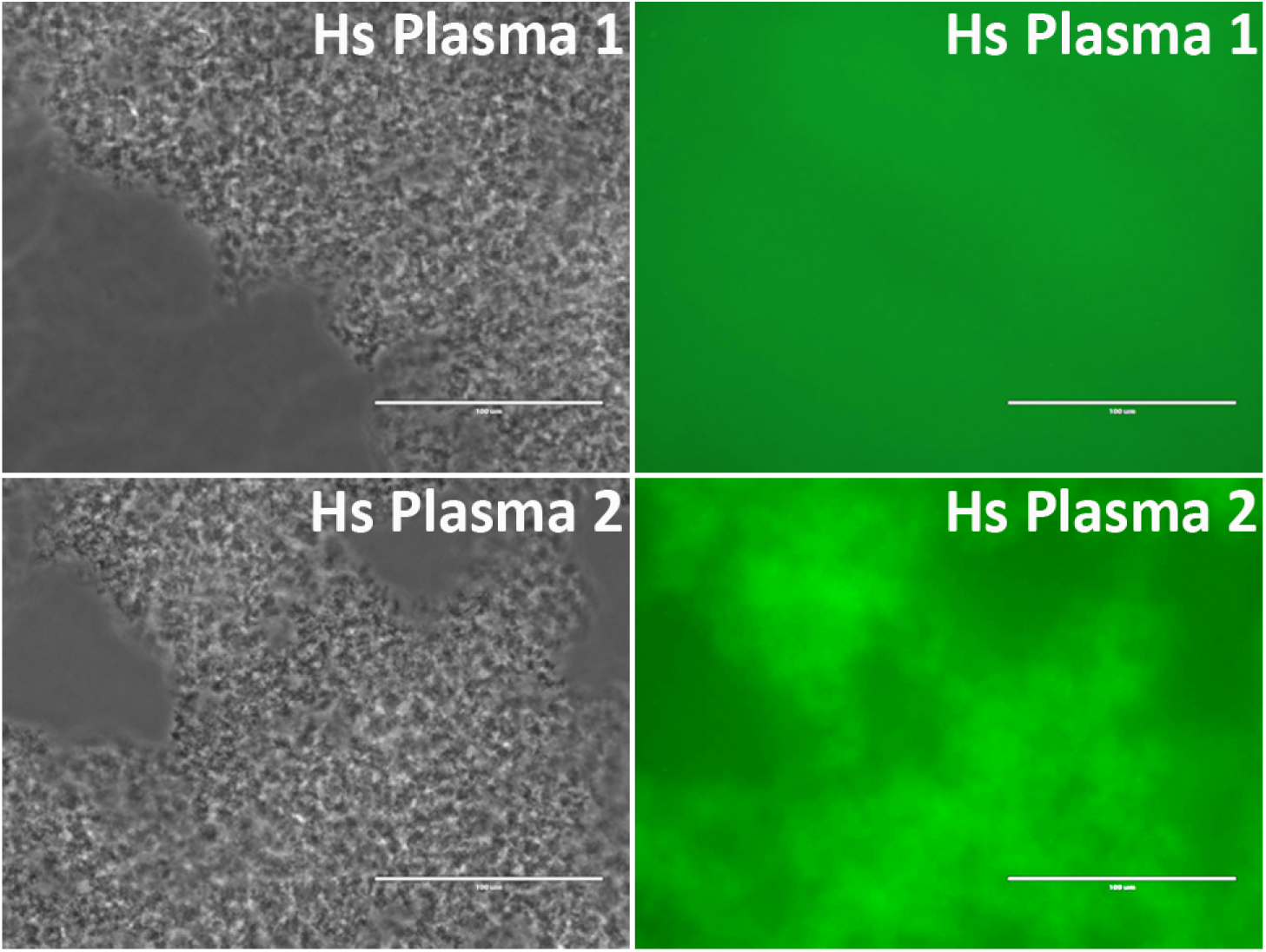
Validation of anti-NMDAR1 autoantibodies with a different NMDAR1-GFP probe. A representative human plasma (Hs Plasma 1) carrying a basal level of anti-NMDAR1 autoantibodies. Human plasma 2 (Hs Plasma 2) carries the highest level of natural anti-NMDAR1 autoantibodies in the 143 human plasma samples. To validate the luciferase-based assay, a GFP-based immunoassay was conducted. Blood antibodies are aggregated by protein A/G/L (gray image). Anti-NMDAR1 autoantibodies within the aggregate from Hs Plasma 2 bind NMDAR1-GFP and emit green fluorescence. Aggregated antibodies from Hs Plasma 1 showed little background.

The anti-NMDAR1 autoantibody-positive plasma identified from the luciferase-based assay were further verified using immunohistochemical staining of mouse hippocampus, the assay recommended for the detection of anti-NMDAR1 autoantibodies in diagnosis of anti-NMDAR encephalitis. NMDAR1 staining signal in hippocampus was stronger from human plasma 2 carrying the highest level of anti-NMDAR1 autoantibodies than from human plasma 1 carrying a basal level of the anti-NMDAR1 autoantibodies **(Figure 5)**. In comparison to the little NMDAR1 staining by control mouse serum **(Figure 2)**, there is some weak NMDAR1 staining from human plasma on mouse hippocampus. This finding is consistent with our cross-species quantification findings that human plasma has a higher basal level of anti-NMDAR1 autoantibody binding activity than mouse serum.

**Figure 5.**
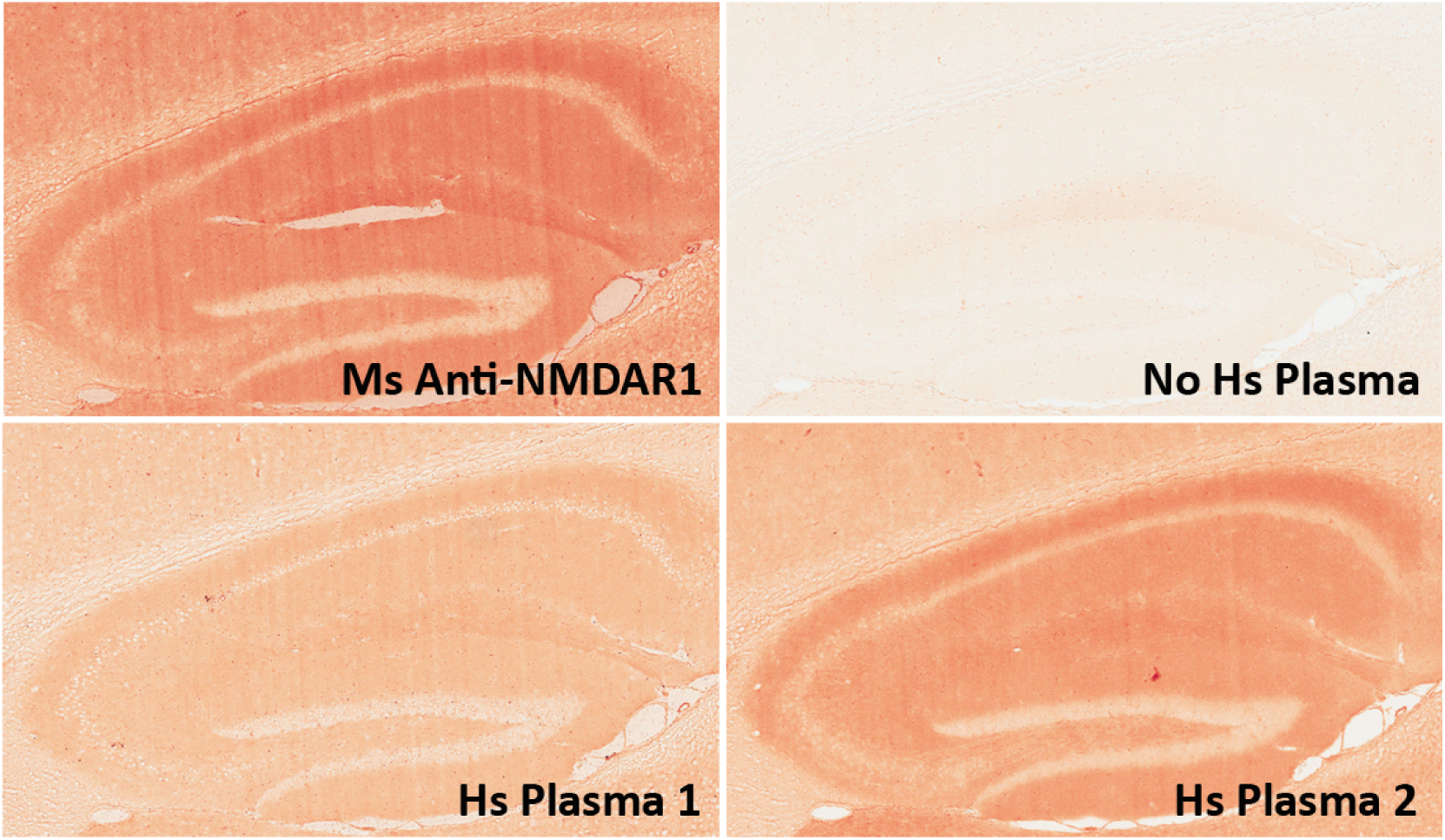
Validation of anti-NMDAR1 autoantibodies with immunohistochemistry. Mouse hippocampal paraffin sections were used for detection of human anti-NMDAR1 autoantibodies. Human plasma was diluted at 1:500 for the IHC analysis. Hippocampal staining with a commercial mouse anti-NMDAR1 monoclonal antibody (1:40,000 dilution) was used as the positive control. A typical hippocampal NMDAR1 staining pattern was observed. Compared to weak signal from Hs Plasma 1 carrying a basal level of anti-NMDAR1 autoantibodies, strong signal was observed from Hs Plasma 2 carrying the highest level of anti-NMDAR1 autoantibodies. A hippocampal section was also included as a negative control without addition of primary antibodies or human plasma but only biotinylated goat anti-human IgG (H + L) secondary antibodies.

Finally, the anti-NMDAR1 autoantibody-positive plasma identified from the luciferase-based assay was verified using the cell-based assay that is commonly used for the detection of anti-NMDAR1 autoantibodies in diagnosis of anti-NMDAR encephalitis. As expected, immunohistochemical staining by human anti-NMDAR1 autoantibodies and mouse anti-NMDAR1 P2 autoantibodies demonstrated complete co-localization **(top panel, Figure 6)**. There are some differential staining intensities between the two different anti-NMDAR1 antibodies. It is possible that the accessibility of different antigenic epitopes of NMDAR1 proteins may contribute to these differences.

**Figure 6.**
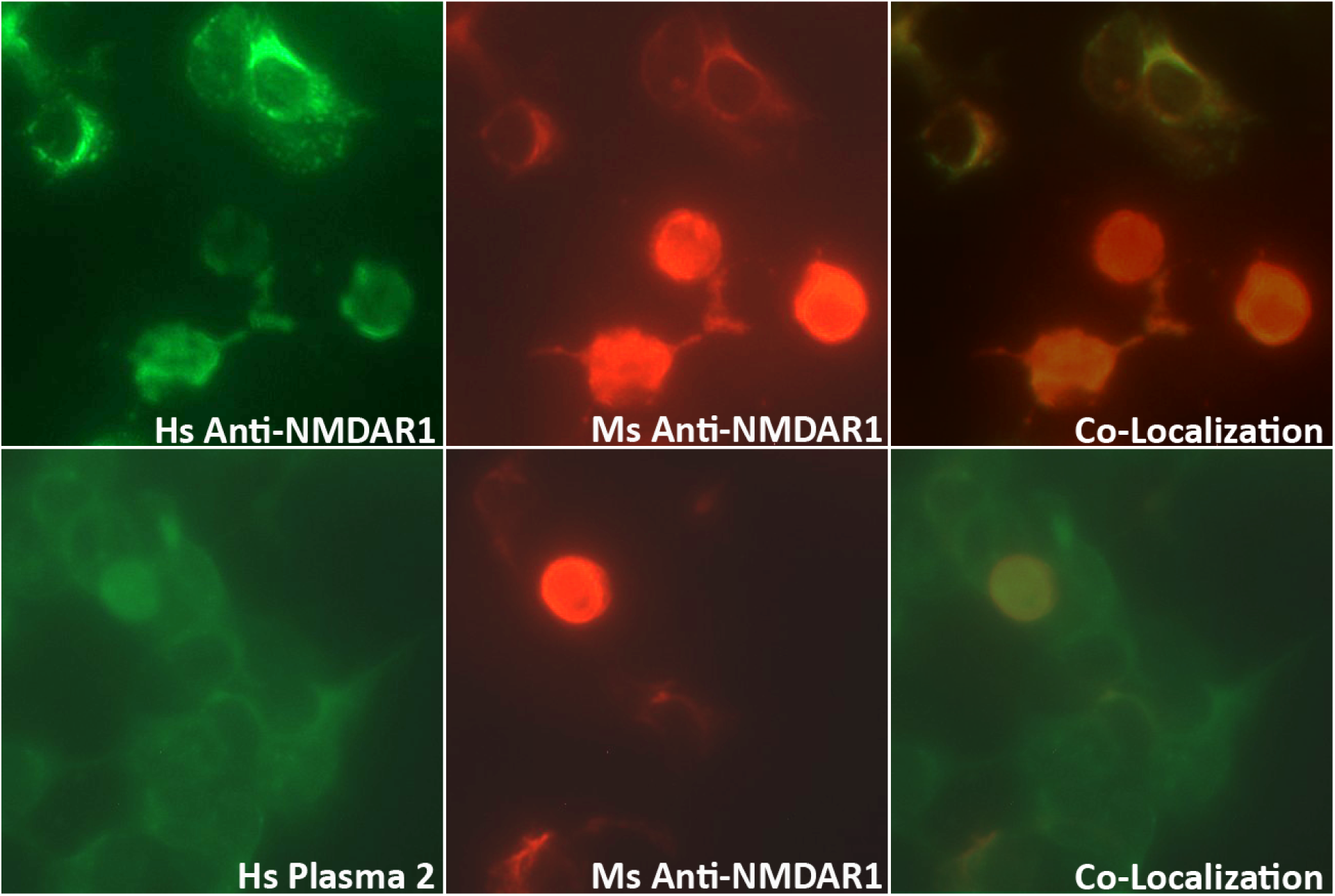
Validation of anti-NMDAR1 autoantibodies with cell-based assay. Human NMDAR1 proteins were expressed on HEK293 cells on BIOCHIPs purchased from Euroimmun. Both anti-Human NMDAR1 autoantibody (Euroimmun) and the mouse anti-NMDAR1 serum (diluted at 1:10) against the NMDAR1 P2 peptide antigens recognize the NMDAR1 proteins on HEK293 cells. The staining between the anti-Human NMDAR1 and mouse anti-NMDAR1 serum was co-localized in cell-based assays (top panel). Both human plasma 2 (diluted at 1:50) and mouse anti-NMDAR1 serum (diluted at 1:10) were used for co-immunofluorescence staining on BIOCHIPS. A high non-specific background was observed for human plasma. Co-localization of the staining between human plasma 2 and mouse anti-NMDAR1 serum (bottom panel), suggesting the presence of human natural anti-NMDAR1 autoantibodies.

To detect lower levels of plasma anti-NMDAR1 autoantibodies found in the general human population, human plasma requires less dilution. However, less diluted plasma generated higher non-specific staining background staining in the cell-based assays **(bottom panel, Figure 6)**. Anti-NMDAR1 autoantibody staining can only be recognized with the help of co-immunocytochemical staining with known anti-NMDAR1 antibodies.

## Discussion

A significant portion of the general human population carry anti-NMDAR1 autoantibodies in their blood circulation, but their effects remain unclear (Zhou, 2021a). In addition, there have been many studies on the potential effects of blood circulating anti-NMDAR1 autoantibodies on psychiatric disorders (Castillo-Gomez et al., 2017; Hammer et al., 2014; Jezequel et al., 2017; Pan et al., 2019), cognitive functions (Yue et al., 2021), and neurodegenerative diseases (Hopfner et al., 2019). All these studies used cell-based assays to detect human blood anti-NMDAR1 autoantibodies and semi-quantify their levels after a series of dilutions. In contrast to high titers of anti-NMDAR1 autoantibodies in patients with anti-NMDAR encephalitis, the levels of anti-NMDAR1 autoantibodies are several orders of magnitude lower in the blood of the general human population and psychiatric patients. High non-specific background staining is inevitable for detection of such lower levels of blood anti-NMDAR1 autoantibodies using this assay. The high background of the cell-based assay is mainly generated from various host cell proteins reacting with different antibodies in human plasma/serum. To minimize this non-specific background, we developed a novel immunoassay for the detection and quantification of blood anti-NMDAR1 autoantibodies, based on our previously reported One-Step immunoassay. In this assay, only antibodies binding NMDAR1 are quantified.

Cell-based assay and immunohistochemical staining are currently the most commonly used methods to quantify blood anti-NMDAR1 autoantibodies. A series of dilutions of serum/plasma are needed for such assays, and detection is subjective. Compared to these labor intensive and less accurate semi-quantification methods, our luciferase-based immunoassay offers an efficient and objective quantification that is particularly helpful for analysis of low levels of blood anti-NMDAR1 autoantibodies in psychiatric patients and the general human population. In addition, our luciferase-based immunoassay can be adapted for high throughput quantification of anti-NMDAR1 autoantibodies when conducting the assay on a large scale. Finally, our quantification method does not require a secondary antibody and protein A/G/L binds antibodies from different mammals, enabling quantification and direct comparison of the levels of blood anti-NMDAR1 autoantibodies across mammalian species.

A potential complication of our luciferase-based immunoassay could arise from antibodies recognizing the *Gaussia* luciferase part of the fusion protein rather than the NMDAR1 antigen part. However, we did not find any human plasma significantly binding the luciferase after screening > 300 human plasma samples. Such false positives, if they occur, can be readily ruled out by cross validating the plasma with NMDAR1-GFP proteins.

Our studies provided proof-of-concept for the development of a simple method to quantify blood anti-NMDAR1 autoantibodies. Different antigens can however be fused with *Gaussia* luciferase for detection of their antibodies after aggregation. The NMDAR1-luciferase fusion proteins can be abundantly produced in *E. coli*, which makes the quantification efficient and cost effective. However, fusion proteins produced from *E. coli* lack post-translational modifications. For detection of antibodies recognizing these modifications, the NMDAR1-luciferase fusion proteins must be produced from mammalian cells. Since protein A/G/L aggregates all isotypes of antibodies, our method cannot directly detect different isotypes of anti-NMDAR1 autoantibodies individually. However, if streptavidin and biotinylated antibodies for a specific isotype of antibodies are used to aggregate antibodies rather than protein A/G/L, it will be feasible to quantify each isotype of blood anti-NMDAR1 autoantibodies.

## Acknowledgements

This work was supported by the grant R01NS135620 (XZ and VR).

## Author contributions

Conceived and designed the experiments: XZ. Performed the experiments: MV, SP, XZ. Analyzed the data: MV, SP, VR, XZ. Wrote the paper: XZ. Edited the paper: MV, SP, VR.

